# Hypersensitivity of the vimentin cytoskeleton to net-charge states and Coulomb repulsion

**DOI:** 10.1101/2024.07.08.602555

**Authors:** Bret A. Unger, Chun Ying Wu, Alexander A. Choi, Changdong He, Ke Xu

## Abstract

As with most intermediate filament systems, the hierarchical self-assembly of vimentin into nonpolar filaments requires no nucleators or energy input. Utilizing a set of live-cell, single-molecule, and super-resolution microscopy tools, here we show that in mammalian cells, the assembly and disassembly of the vimentin cytoskeleton is highly sensitive to the protein net charge state. Starting with the intriguing observation that the vimentin cytoskeleton fully disassembles under hypotonic stress yet reassembles within seconds upon osmotic pressure recovery, we pinpoint ionic strength as its underlying driving factor. Further modulating the pH and expressing differently charged constructs, we converge on a model in which the vimentin cytoskeleton is destabilized by Coulomb repulsion when its mass-accumulated negative charges (-18 per vimentin protein) along the filament are less screened or otherwise intensified, and stabilized when the charges are better screened or otherwise reduced. Generalizing this model to other intermediate filaments, we further show that whereas the negatively charged GFAP cytoskeleton is similarly subject to fast disassembly under hypotonic stress, the cytokeratin, as a copolymer of negatively and positively charged subunits, does not exhibit this behavior. Thus, in cells containing both vimentin and keratin cytoskeletons, hypotonic stress disassembles the former but not the latter. Together, our results both provide new handles for modulating cell behavior and call for new attention to the effects of net charges in intracellular protein interactions.

## Introduction

Complex cellular structures are sometimes assembled from simple components. The vimentin cytoskeleton, the predominant intermediate filament system of mesenchymal cells, is a homopolymer hierarchically self-assembled from a single 54 kDa protein. It plays key roles in cell mechanics, structural integrity, and beyond, and is a marker in epithelial-mesenchymal transition (*1-7*). Whereas the vimentin cytoskeleton is mechanically strong and highly resistant to stress, recent studies have reported its unexpected fast disassembly and reconfiguration under hypotonic (low osmotic pressure) conditions (*8, 9*), pointing to potentially new mechanisms of its structural dynamics. More generally, as intermediate filaments are typically self-assembled from monomers in a nonpolar fashion requiring no nucleators or energy input (*1-4, 7, 10*), it remains unclear what factors drive their assembly and disassembly processes in the cell.

Utilizing a set of live-cell, single-molecule, and super-resolution microscopy tools, including stochastic optical reconstruction microscopy (STORM) (*11-13*) and single-molecule displacement/diffusivity mapping (SM*d*M) (*14*), here we show that in the mammalian cell, the hypotonic-stress-disassembled vimentin cytoskeleton reassembles into filaments within seconds upon osmotic pressure recovery. By varying the medium ionic strength and pH, as well as expressing vimentin constructs with differently charged linkers, we next elucidate a novel mechanism in which the stability of the vimentin cytoskeleton is dictated by the protein net charges, so that the crowded accumulation of net negative charges (∼−18 per vimentin protein) in the filament gives rise to Coulomb repulsion and instability. Generalizing this model to other intermediate filament systems, we further show that whereas cytoskeleton of the similarly negatively charged glial fibrillary acidic protein (GFAP) is also subject to fast dissociation under hypotonic stress, the cytokeratin, as a copolymer of negatively and positively charged subunits, does not exhibit this behavior. Together, our results both highlight the fundamental importance of net charges in intracellular protein interactions and provide new handles for modulating cell structure and behavior.

## Results

We start by confirming previous observations (*8, 9*). With fixed and immunostained COS-7 cells, we thus reaffirmed that 5-min hypotonic treatments with water led to complete disassembly of the vimentin cytoskeleton, with STORM super-resolution microscopy resolving no remaining filaments at its ∼20 nm resolution (**Fig. S1**). However, we further found that for cells that underwent the same hypotonic treatment but then recovered in the isotonic cell medium for 30 min, the vimentin cytoskeleton fully reassembled, with filaments extending throughout the cell (**Fig. S1**).

To monitor the dynamics of the above process, we expressed in COS-7 cells vimentin tagged by the fluorescent protein (FP) mEos3.2. Live-cell fluorescence microscopy (**Fig. 1A**) thus showed that after hypotonic treatment with water, the vimentin cytoskeleton quickly disassembled in ∼2 min (**Fig. 1A ii**) and became fully dispersed in the cytoplasm by 3 min (**Fig. 1A iii**). Remarkably, upon next reverting to the isotonic medium, reassembly occurred immediately. Within a few seconds, the vimentin fluorescence condensed into fibrils-like foci, leaving little diffusive signal in the cytoplasm (**Fig. 1A iv**). These fibrils then progressively joined into fibers and continued to develop into longer filaments over minutes (**Fig. 1A v**). Extensive filaments were thus established in the cell by 15 min (**Fig. 1A vi**).

**Figure 1.**
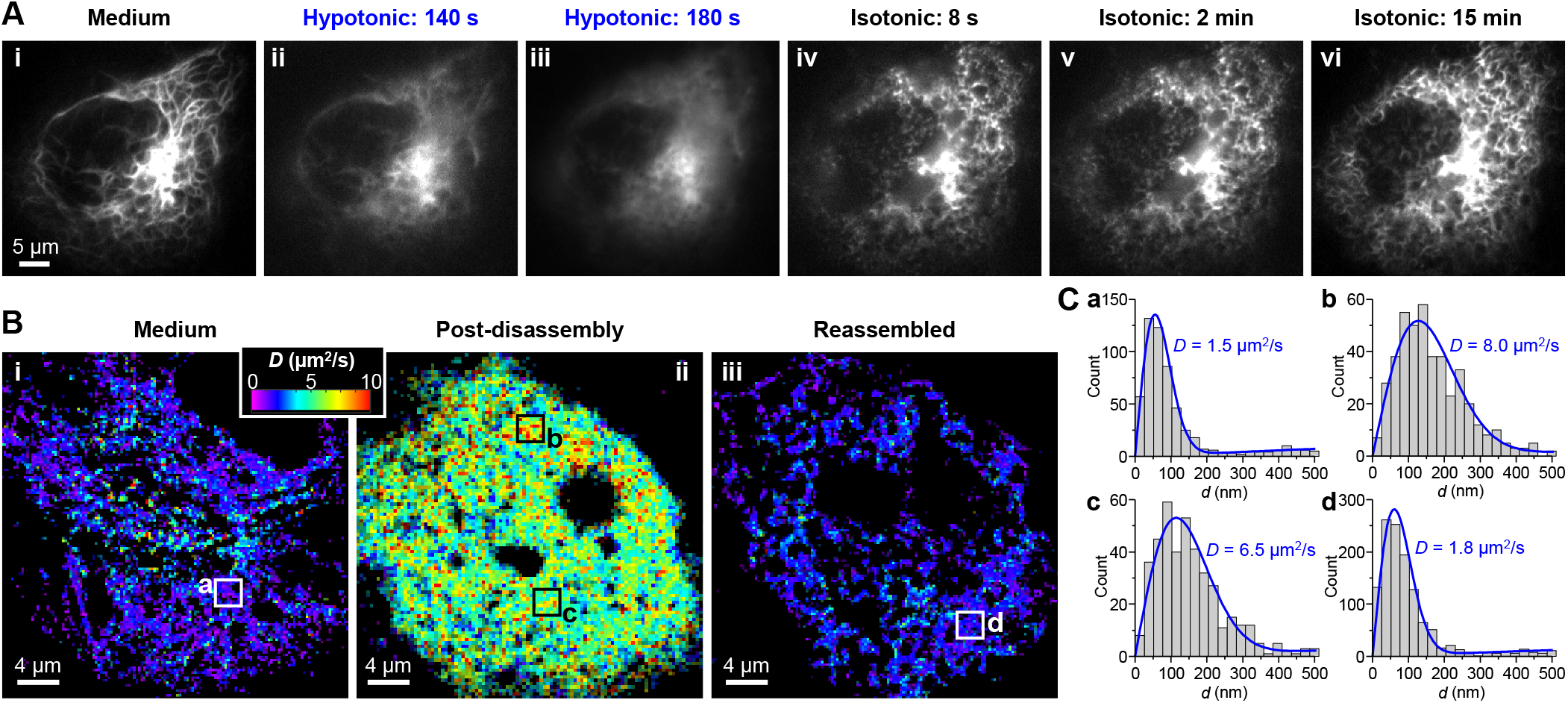
Live-cell fluorescence microscopy and SM*d*M unveil fast disassembly and even faster reassembly of the vimentin cytoskeleton under osmotic pressure drop and recovery. (**A**) Fluorescence micrographs of vimentin-mEos3.2 expressed in a living COS-7 cell, (**i**) initially in an isotonic cell medium, (**ii-iii**) under hypotonic stress in water for 140 s (**ii**) and 180 s (**iii**), and then (**iv-vi**) after returning to the isotonic medium for 8 s (**iv**), 2 min (**v**), and 15 min (**vi**). (**B**) Color-coded SM*d*M maps of the local diffusion coefficient *D* for vimentin-mEos3.2 expressed in a living COS-7 cell, (**i**) initially in an isotonic medium, (**ii**) under hypotonic treatment for 5-22 min, so that the vimentin cytoskeleton had disassembled, and (**iii**) after next reverting to the isotonic medium for 0.5-5 min, so that the vimentin cytoskeleton had reassembled. (**C**) Distribution of the SM*d*M-measured displacements of single vimentin-mEos3.2 molecules in 1 ms time windows, for the boxed regions marked as a-d in (B). Blue curves: Fits to the SM*d*M diffusion model, with resultant *D* values marked in each plot.

To further elucidate the vimentin assembly state in the disassembly-reassembly process, we utilized SM*d*M, a single-molecule super-resolution tool we recently developed that uniquely maps intracellular diffusion with high fidelity and spatial resolutions (*14*). Whereas our previous live-cell applications of SM*d*M have focused on the diffusion patterns of untagged fluorescent tracers (*14-16*), here we repurpose SM*d*M to assess the intracellular assembly state of vimentin-mEos3.2. Through repeated execution of paired stroboscopic excitation pulses across tandem camera frames (*14*), we thus collected the transient (1 ms) nanoscale displacements of ∼10^5^ individual vimentin-mEos3.2 molecules in a COS-7 cell as we imposed and withdrew hypotonic stresses. Spatially binning the accumulated single-molecule displacements for local statistics (*14, 17*) next yielded spatial maps of the diffusion coefficient *D*.

In the untreated cell, we thus found that in the filament state, vimentin-mEos3.2 exhibited limited motion with apparent *D* ∼1.5 µm^2^/s (**Fig. 1B i** and **Fig. 1C a**). After hypotonic disassembly, vimentin-mEos3.2, now redistributed homogenously in the cytoplasm, showed substantially increased *D* of ∼6-8 µm^2^/s (**Fig. 1B ii** and **Fig. 1C bc**). These values are ∼1/3 of free mEos3.2 FP in mammalian cells (*14*). Whereas vimentin-mEos3.2 is ∼3-fold heavier than mEos3.2, *D* scales inversely to the cubic root of molecular weight with little dependence on the protein shape (*18-20*). The ∼6-8 µm^2^/s value of vimentin-mEos3.2 thus translates to an average oligomer size of ∼10 vimentin. This result is consistent with the STORM observation of immunolabeled fixed cells, in which no filament fragments remained after the disassembly (**Fig. S1**). After osmotic recovery, the SM*d*M-determined *D* rapidly dropped back to <2 µm^2^/s throughout the cell as vimentin reassembled into short fibers (**Fig. 1B iii** and **Fig. 1C d**).

Together, we have shown that while the vimentin cytoskeleton disassembles rapidly under hypotonic stress, it reassembles even faster upon osmotic-pressure recovery. Previous studies have attributed hypotonicity-induced vimentin cytoskeleton disassembly and rearrangements to microtubule-based transport (*8*) or calcium-activated proteolysis (*9*). The prompt reassembly of vimentin filaments we unveiled in osmotic-pressure recovery does not fit either model. Consequently, we seek an alternative mechanism that more directly modulates vimentin’s properties. Osmotic shifts drive water into or out of the cell. Resultant dilution or concentration of cytoplasmic contents have been discussed in terms of changes in macromolecular crowding and viscosity (*21-24*), including the recently reported controls of intracellular microtubule polymerization (*24*) and protein phase separation (*23*). We reason, however, that the unexpectedly fast vimentin disassembly-reassembly above may instead be driven by changes in intracellular ionic strength (*25*).

The *in vitro* polymerization of vimentin is typically initiated by adding salts (*26-30*), requiring no nucleators or energy input. Recent studies further reported *in vitro* modulations of the vimentin filament structure and stability with varied ionic strengths and pH (*30, 31*), the addition of multivalent cations (*32, 33*), and the modification of vimentin net charge (*34*). These results are generally consistent with a model in which, as the vimentin filament is densely assembled from a 54 kDa protein that carries an ∼−18 net charge at the physiological pH of ∼7.3 (**Table S1** and **Fig. 2D** below), Coulomb repulsion may destabilize the filament when charges are not well screened (**Fig. 2C**). However, it is unclear if this mechanism could be relevant in the cell. Although net charge-driven protein interactions in the mammalian cell have gained recent attention, the focus has been on attractions between opposite charge signs (*14, 16, 35*) and spatial exclusions due to the same charge sign (*36*).

**Figure 2.**
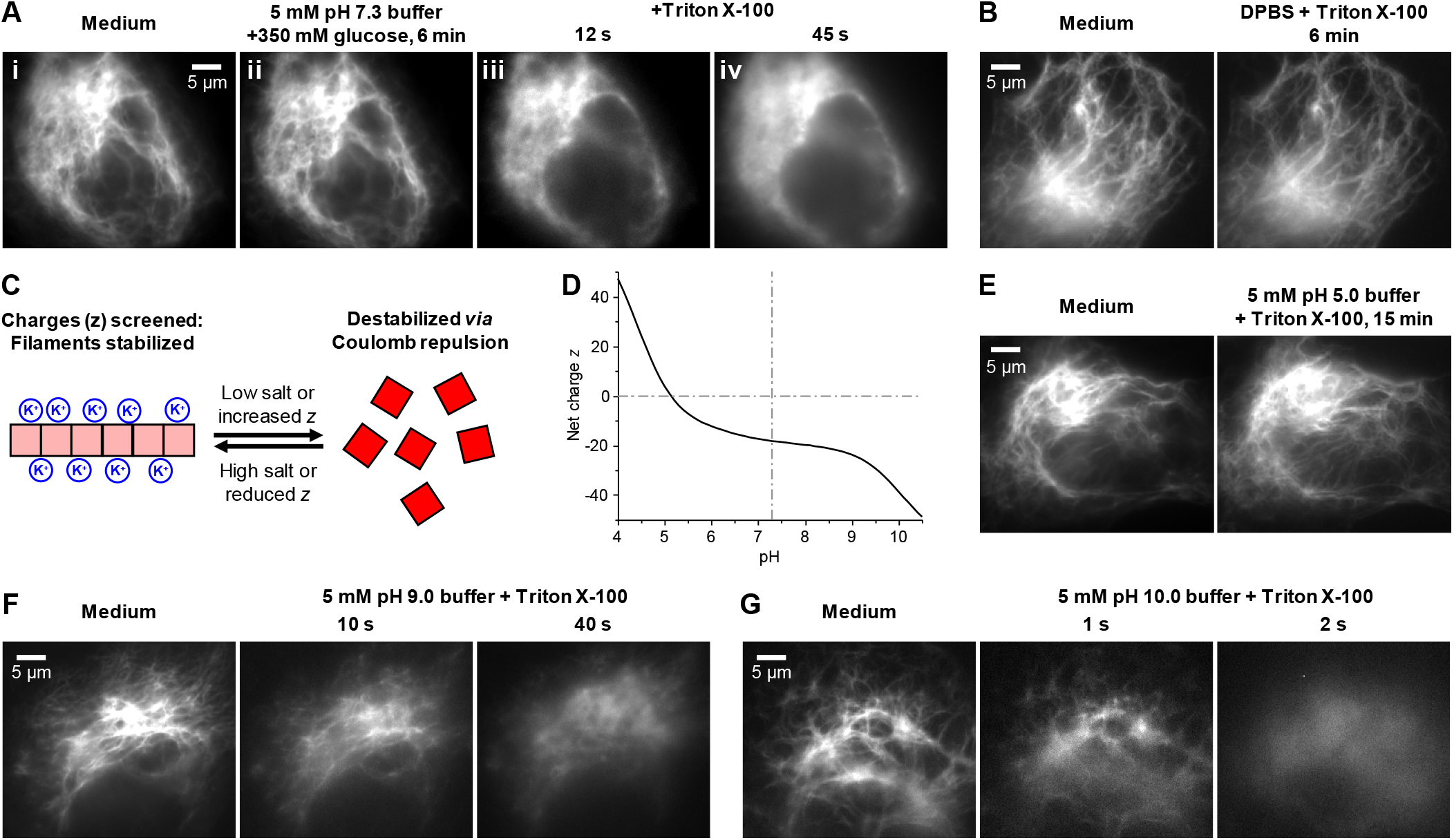
Fluorescence microscopy of vimentin-mEos3.2 in COS-7 cells permeabilized under different ionic strengths and pHs unveils hypersensitivity of the vimentin cytoskeleton stability to protein net charges. (**A**) Fluorescence micrographs of a cell (**i**) initially in an isotonic cell medium, (**ii**) 6 min after replacing the medium with a 5 mM phosphate buffer (pH = 7.3) with 350 mM glucose added, and (**iii**,**iv**) 12 s and 45 s after next adding 0.2% Triton X-100 into the medium. (**B**) Fluorescence micrographs for another cell before and 6 min after changing the cell medium to DPBS with 0.2% Triton X-100. (**C**) Model: The vimentin filament is stabilized when the accumulated negative net charges (red) are ionically screened or otherwise reduced, and destabilized when the charges are less screened or otherwise intensified. (**D**) The expected net charge of vimentin as a function of pH, estimated with Protein Calculator v3.4 (http://protcalc.sourceforge.net). (**E**) Fluorescence micrographs for a cell before and 15 min after changing the cell medium to a 5 mM acetate buffer (pH = 5.0) with 0.2% Triton X-100. (**F**) Fluorescence micrographs for another cell before and after replacing the medium with a 5 mM CAPSO buffer (pH = 9.0) with 0.2% Triton X-100 at 10 s and 40 s. (**G**) Fluorescence micrographs for another cell before and after replacing the cell medium with a 5 mM CASPO buffer (pH = 10.0) with 0.2% Triton X-100 at 1 s and 2 s.

We set out to first decouple the effects of ionic strength and osmotic pressure. To this end, we first replaced the cell medium with a low ionic-strength but high osmotic-pressure medium, a 5 mM phosphate buffer (pH = 7.3) with 350 mM glucose added. As the added glucose upshifted the osmolarity, cells did not experience hypotonic stress, and the vimentin cytoskeleton remained undisturbed for 6 min (**Fig. 2A i,ii**). Next permeabilizing the cell membrane with Triton X-100, which equilibrated the cytosol with the medium, led to fast disassembly of the vimentin cytoskeleton in 45 s (**Fig. 2A iii,iv**). This result was independent of how the osmolarity or permeabilization was achieved, as we observed similar results when sorbitol and saponin were used instead (**Fig. S2**). In comparison, for cells permeabilized in Dulbecco’s Phosphate-Buffered Saline (DPBS), which matches the cytosol ionic strength, the vimentin cytoskeleton remained intact (**Fig. 2B**). These results suggest that lowered ionic strength, rather than osmolarity, drives the disassembly of the vimentin cytoskeleton, consistent with our model that the densely accumulated negative charges may destabilize vimentin filaments when not sufficiently screened (**Fig. 2C**).

To further probe the sensitivity of the vimentin cytoskeleton stability to the protein charge state, we varied the medium pH (**Fig. 2D**). Notably, as we permeabilized cells in a 5 mM acetate buffer at pH = 5.0 to match the isoelectric point of vimentin (**Fig. 2D**), the vimentin cytoskeleton remained intact in this low ionic-strength medium (**Fig. 2E**). In the opposite direction, for cells permeabilized in 5 mM CAPSO buffers at pH = 9.0 and 10.0, progressively faster vimentin disassembling was observed (**Fig. 2FG**) versus the pH = 7.3 phosphate buffer as vimentin carried higher negative net charges. Notably, by more-than-doubling the vimentin net charge (**Fig. 2D**), the pH = 10.0 condition led to instantaneous filament disassembly within 2 s (**Fig. 2G**). Increasing the ionic strengths of the high-pH media to physiological levels impeded but did not prevent disassembly (**Fig. S2**). Together, these results showed that the vimentin cytoskeleton stability is highly sensitive to the protein charge state, and so unscreened high net charges induce filament disassembly (**Fig. 2C**).

To further examine the net-charge effects under physiological pH, we modified the net charge on the expressed vimentin-mEos3.2 and returned to assess the stability of the cytoskeleton under hypotonic stresses without permeabilizing the cell. Whereas mEos3.2 is ∼0-charged, we introduced differently charged 11 amino-acid linkers between the vimentin and mEos3.2 sequences to shift the total net charge of the expressed proteins (**Table S1**) (*14, 16*). We thus found that with the expression of vimentin-(−5)-mEos3.2 containing a (−5)-charged linker, hypotonic stress-induced vimentin disassembly was accelerated, so that fast disassembly was noted after ∼60 s and vimentin was homogenized in the cytoplasm by 120 s (**Fig. 3A**). Meanwhile, with the expression of vimentin-(+6)-mEos3.2 containing a (+6)-charged linker, hypotonic stress-induced vimentin disassembly was substantially impeded, which occurred slowly after ∼300 s (**Fig. 3B**). Thus, increasing or reducing the total negative charges by introducing extraneous negative or positive charges respectively destabilized and stabilized the vimentin cytoskeleton, further supporting our model (**Fig. 2C**).

**Figure 3.**
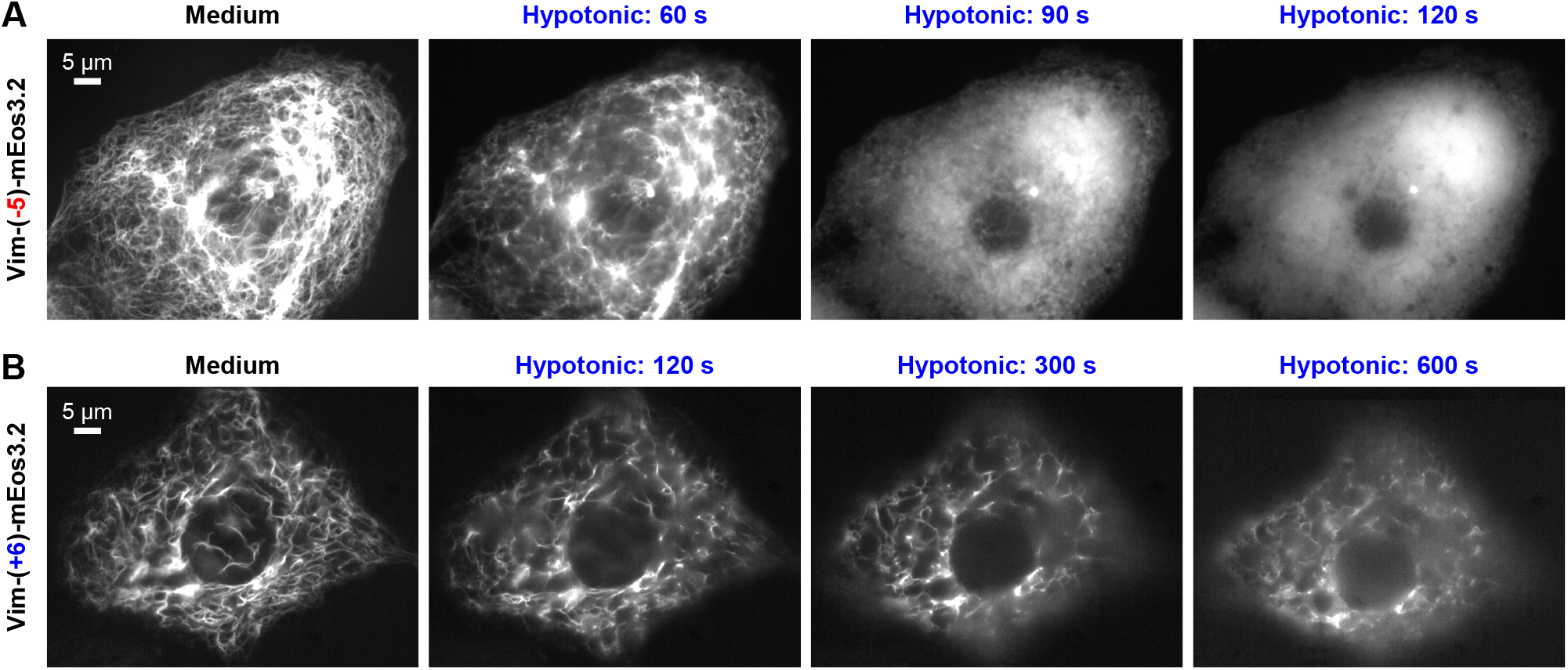
Live-cell fluorescence microscopy of COS-7 cells expressing vimentin-mEos3.2 constructs with differently charged linkers further underscores protein net charge as a key factor in vimentin cytoskeleton stability. (**A**) Representative fluorescence micrographs with vimentin-(-5)-mEos3.2, in which a (-5)-charged 11 amino-acid linker was inserted between vimentin and mEos3.2 sequences, after hypotonic treatment (5 mM phosphate buffer, pH = 7.3) of different durations. (**B**) Representative fluorescence micrographs with vimentin-(+6)-mEos3.2, in which a (+6)-charged 11 amino-acid linker was inserted between vimentin and mEos3.2 sequences, after the same hypotonic treatment of different durations.

We next generalize the above net-charged-based mechanisms to other intermediate-filament systems. Cytokeratin constitutes the characteristic intermediate filament in epithelial cells (*37-39*). Contrasting with the single-component vimentin homopolymer, the cytokeratin is a copolymer of Type I and Type II keratins, which form an obligatory heterodimer before subsequent assembly into tetramers and filaments. Whereas Type I keratins are acidic, Type II keratins are basic or near-neutral (*37-39*). We reason that as the keratin filament is thus assembled from a combination of subunits with negative, positive, and near-neutral charges, it should not accumulate significant Columb repulsions and so would not be destabilized when challenged by low ionic-strength conditions, *e*.*g*., those due to hypotonic treatments.

We start by examining untransfected PtK2 cells, in which endogenous vimentin and keratin intermediate filaments coexist and interact (*40*). We thus found that for PtK2 cells fixed after 5 min hypotonic treatment in a 5 mM phosphate buffer (pH = 7.3), cytokeratin (as immunostained with a pan-cytokeratin antibody) retained filament structures (**Fig. 4A i**), even as vimentin cytoskeleton in the same cells fully disassembled (**Fig. 4A ii,iii**). STORM showed diffuse vimentin patterns with no discernible filaments (**Fig. 4A iv**), similar to results in COS-7 cells.

**Figure 4.**
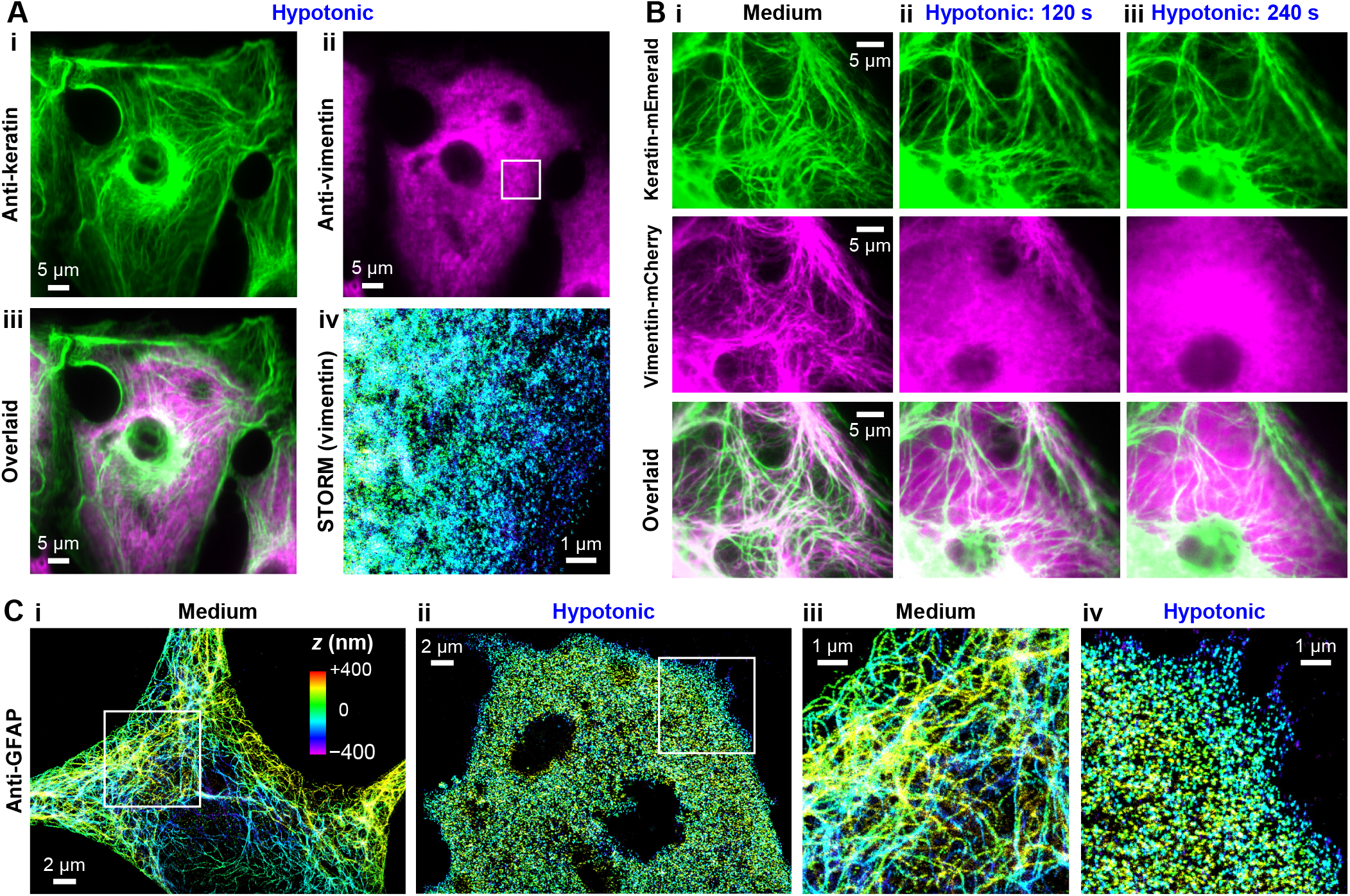
Comparison with cytokeratin and GFAP further generalizes the net-charge mechanism of intermediate-filament intracellular stability. (**A**) Comparison of immunofluorescence micrographs of endogenous cytokeratin and vimentin in fixed wild-type PtK2 cells subjected to hypotonic treatment of 5 mM phosphate buffer (pH = 7.3) for 5 min. (**i**) Anti-pan cytokeratin, (**ii**) Anti-vimentin, (**iii**) Overlaid image, (**iv**) 3D-STORM super-resolution image of anti-vimentin for the boxed region in (ii). (**B**) Two-color live-cell fluorescence microscopy of a PtK2 cell co-transfected with keratin-mEmerald and vimentin-mCherry, shown as separate and merged images, before (**i**) and after (**ii**,**iii**) hypotonic treatment in a 5 mM phosphate buffer for 120 s (**ii**) and 240 s (**iii**). (**C**) 3D-STORM super-resolution images of anti-GFAP for primary astrocytes isolated from the rat hippocampus, for samples without (**i**) and with (**ii**) hypotonic treatments in a 5 mM phosphate buffer for 5 min. (**iii**,**iv**) Zoom-in of the boxes in (i,ii). Colors in (A iv) and (C) encode axial (depth) position, based on the color scale shown in (C i).

To monitor the above process, we next performed two-color live-cell imaging for PtK2 cells co-expressing vimentin and keratin tagged by two different FPs. We thus found that in the untreated cells, the two intermediate filament systems partially colocalized and interweaved (**Fig. 4B i**). Upon hypotonic treatment, the vimentin filaments quickly disassembled in ∼2 min (**Fig. 4B ii**) and became fully diffusive in the cell by 4 min (**Fig. 4B iii**). Over the same period, no significant structural changes were observed for the keratin filaments, although cell swelling dislocated some filaments and shifted some out of focus (**Fig. 4B** and **Movie S1**). Thus, even though initially colocalized and likely mechanically coupled, vimentin but not keratin filaments disassembled under hypotonic stress.

We next turned to glial fibrillary acidic protein (GFAP) (*4, 41*). As implied by its name, GFAP is acidic, and it carries ∼−12 net charges at the physiological pH (**Table S1**). In astrocytes, GFAP polymerizes into intermediate filaments in a fashion analogous to vimentin. With primary rat hippocampus astrocytes, we thus showed full disassembly of the endogenous GFAP cytoskeleton under hypotonic stresses, with STORM super-resolution microscopy resolving no remaining filaments (**Fig. 4C**).

Together, a comparison between three intermediate filament systems generalized our model, in which the excessive same net charges accumulated in the densely packed vimentin and GFAP filaments give rise to Coulomb repulsion and instability under low ionic strengths, whereas the cytokeratin filament is unsusceptible to such effects due to its mixed charge signs of Type I and II subunits.

## Discussion

With the surprising discovery of ultrafast disassembly and reassembly of the vimentin cytoskeleton in osmotic-pressure drop and recovery, in this work we formulated a model in which the concentrated negative net charges on the assembled filaments, when not ionically screened or otherwise abated, cause structural instability. By decoupling the effects of ionic strength and osmotic pressure with added osmolytes and further varying the pH, our results on cells permeabilized under different conditions supported this new model. Hypotonic assays of cells expressing vimentin with differently charged linkers further highlighted the sensitivity of the vimentin cytoskeleton to protein charge states. A comparison with cytokeratin and GFAP next generalized the above net-charge-based mechanisms to other intermediate filament systems.

The intermediate filaments are hierarchically assembled from monomers in a nonpolar fashion. The self-assembly process requires no nucleators or energy input, and it remains unclear what factors drive their assembly and disassembly in the cell. The vimentin filament is densely assembled from a single protein of high negative net charge, and *in vitro* experiments generally indicate that the vimentin filament formation and mechanical strength are promoted when the accumulated negative net charges are ionically screened or otherwise reduced, and *vice versa*. Here we showed that in the mammalian cell, protein net charge plays a key role in the fast disassembly and reassembly of the vimentin cytoskeleton in response to changes in ionic strength and pH. Whereas among these experiments we artificially altered protein net charges by adding charged linkers, for endogenous vimentin, enhanced accumulation of negative charges may be achieved *via* phosphorylation. Indeed, phosphorylation generally weakens intermediate filaments and promotes disassembly (*34, 42, 43*). The different factors may synergistically modulate the intermediate filament charge states to induce fast cytoskeletal reconfiguration. For example, during mitosis, as phosphorylation upregulation (*44-46*) steers the vimentin cytoskeleton toward instability, the transient, up to ∼30% volume increase and recovery (*47-50*) may help drive the necessary vimentin disassembly and rearrangement (*51, 52*) through changes in intracellular ionic strengths.

Meanwhile, whereas our results on GFAP gave another example of Coulomb repulsion-induced filament instability, our comparison with cytokeratin highlighted behavior differences for intermediate filaments assembled from a mixture of differently charged subunits. In cells containing both vimentin and keratin filaments, we thus showed that hypotonic treatments disassembled the former but not the latter, further underscoring Coulomb repulsion between the same charge sign in filament destabilization. Notably, the gradual changeover from keratin to vimentin intermediate-filament systems is a key trait of epithelial-mesenchymal transition, and the coexistence of both systems has been suggested as an indicator for aggressive cancer cells (*6, 53*). Our finding that hypotonic stress selectively disassembles the vimentin but not the keratin cytoskeleton provides a potentially powerful handle for controlling cell behaviors in this context.

As we have successfully explained the contrasting behaviors of three intermediate filament systems with our new model, future studies could extend our model to understand the stability of other densely assembled cellular structures. More fundamentally, together with recent studies highlighting intracellular attractions due to opposite charge signs and spatial exclusions due to the same charge sign (*14, 16, 35, 36*), our results call for new attention to the effects of net charges in intracellular protein interactions.

## Acknowledgment

We acknowledge support by the National Institute of General Medical Sciences of the National Institutes of Health (R35GM149349), the Packard Fellowships for Science and Engineering, the Pew Charitable Trusts, and the Heising-Simons Faculty Fellows Award.

## Supporting Information

### Material and methods

#### Cell culturing

COS-7 and PtK2 cells (University of California Berkeley Cell Culture Facility) were cultured in Dulbecco’s Modified Eagle’s Medium (DMEM, high glucose) with the addition of 10% fetal bovine serum (FBS), 1× non-essential amino acids (NEAA), and 1× GlutaMAX Supplement (for PtK2 cells only), in 5% CO_2_ at 37 ^°^C. For live-cell and SM*d*M imaging, cells were plated in 8-well chambered coverglasses (Nunc Lab-Tek II). For STORM imaging, cells were plated on 18 mm #1.5 glass coverslips. Primary rat hippocampus astrocytes were from BrainBits LLC, and were cultured in the NbAstro medium (BrainBits) and plated on coverslips coated with poly-D-lysine, per recommended protocols.

#### Plasmid constructs and transfection

Vimentin-mEos3.2 (Addgene #57485), vimentin-mCherry (Addgene #55156), and keratin-mEmerald (KRT18-mEmerald; Addgene #54134) plasmids were gifts from Michael Davidson. Vimentin-(-5)-mEos3.2 and vimentin-(+6)-mEos3.2 (**Table S1**) were prepared by first cutting out the mEos3.2 from Vimentin-mEos3.2 with BamHI and NotI restriction enzymes. The plasmids were then ligated through Gibson Assembly with the PCR product of vimentin-mEos3.2 and the desired linkers. The constructed plasmids were amplified in XL1-Blue cells and extracted with the QIAprep Spin Miniprep Kit (QIAGEN). The DNA sequences were verified by Sanger sequencing at the UC Berkeley DNA Sequencing Facility. Cells were transfected with the Neon Transfection System (ThermoFisher) or Lipofectamine 3000 (ThermoFisher) according to the recommended protocols, 48-72 hours before imaging.

#### Live-cell imaging

Unless otherwise mentioned, for live-cell imaging, the cell culture medium was first replaced with an isotonic imaging buffer (*14*), Leibovitz’s L-15 medium (Gibco 21083027) supplemented with 20 mM HEPES (pH 7.3; Gibco 15630106). Dulbecco’s phosphate-buffered saline (DPBS) was from Gibco (14040133). The 5 mM pH = 7.3 phosphate buffer was prepared by diluting a 0.1 M phosphate buffer (Sigma-Aldrich, P5244). The 5 mM pH = 5.0 acetate buffer was prepared by diluting a 0.1 M potassium acetate buffer (Sigma-Aldrich, SAE0157). The pH = 9.0 and 10.0 CAPSO buffers were prepared by mixing the CAPSO acid (Sigma-Aldrich C2278) and sodium salt (Sigma-Aldrich C2154) at different ratios and diluting to 5 mM. Triton X-100, saponin, glucose, sorbitol, and KCl (Sigma-Aldrich) were added to the described amounts. Epifluorescence imaging was performed on a Nikon Ti-E inverted fluorescence microscope with an oil-immersion objective (Nikon CFI Plan Apochromat λ 100×, NA 1.45) or an Olympus IX73 inverted epifluorescence microscope with a water-immersion objective (Olympus, UPLSAPO60XW, NA 1.2).

#### SM*d*M of live cells

SM*d*M was performed on a setup based on a Nikon Ti-E inverted fluorescence microscope, as described previously (*14, 15*). Briefly, excitation and photoactivation lasers at 561 and 405 nm were focused at the back focal plane of an oil-immersion objective lens (Nikon CFI Plan Apochromat λ 100×, NA 1.45) toward the edge of the objective lens to illuminate a few micrometers into the cell. Single-molecule images were continuously recorded in the wide field with an EM-CCD (iXon Ultra 897, Andor) at a framerate of 110 Hz. The 561 nm excitation laser was repeatedly applied as tandem excitation pulses of *τ* = 500 µs duration across paired camera frames with a center-to-center separation of Δ*t* = 1 ms, so that single-molecule displacements were recorded across the paired frames for the fixed 1 ms time window. For analysis, the single-molecule displacements accumulated over a given period were first spatially binned onto a 240×240 nm^2^ grid. For each spatial bin, the accumulated single-molecule displacements were fitted through maximum likelihood estimation (MLE) to a modified two-dimensional random walk model with the probability distribution 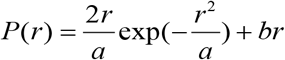, where *r* is the single-molecule displacement, *a* = 4*D*Δ*t, D* being the diffusion coefficient, and *b* accounts for the background. By thus determining the *D* values for each spatial bin, color-coded SM*d*M maps of local *D* were generated.

#### Immunofluorescence and 3D-STORM of fixed cells

Three-dimensional stochastic optical reconstruction microscopy (3D-STORM) was performed for fixed cells as described previously (*11, 12, 54*). Briefly, cells were fixed with 3% (w/v) paraformaldehyde and 0.1% (w/v) glutaraldehyde in DPBS for 20 min. After reduction with a freshly prepared 0.1% sodium borohydride solution in PBS for 5 min, the sample was permeabilized and blocked in a blocking buffer (3% w/v bovine serum albumin (BSA) and 0.1% v/v Triton X-100 in DPBS) for 1 h. Afterward, the cells were incubated with primary antibodies in the blocking buffer for 12 h at 4 ^°^C. After washing in a washing buffer (0.3% w/v BSA and 0.01% v/v Triton X-100 in DPBS) three times, the cells were incubated with dye-labeled secondary antibodies for 1 h at room temperature. Then, the samples were washed three times with the washing buffer and three times with PBS. Primary antibodies used: rabbit anti-vimentin (Cell Signaling Technology, 5741, 1:100), rabbit anti-GFAP (Proteintech, 16825-1-AP, 1:200), and mouse anti-pan cytokeratin (Sigma-Aldrich, C2562, 1:100). Secondary antibodies used: Alexa Fluor 647-labeled goat anti-rabbit (Invitrogen, A21245) for STORM imaging and donkey anti-mouse (Jackson ImmunoResearch, 715-005-151) conjugated with CF568 succinimidyl ester (Biotium, 92131) for epifluorescence imaging in a second color channel when needed. The immunolabeled sample was imaged in a Tris-Cl buffer (pH 7.5) containing 100 mM cysteamine, 5% glucose, 0.8 mg/mL glucose oxidase, and 40 µg/mL catalase. The 647 nm laser illuminated the sample at ∼2 kW/cm^2^, which photoswitched most of the labeled dye molecules into a dark state while allowing a small, random fraction of molecules to emit across the wide field over different camera frames. Single-molecule emission was passed through a cylindrical lens of focal length 1 m to introduce astigmatism, and recorded with an Andor iXon Ultra 897 EM-CCD camera at a framerate of 110 Hz. A total of ∼50,000 frames were recorded for each STORM run. The recorded single-molecule images were localized and rendered as super-resolution images as described previously (*11, 12*).

**Table S1.**
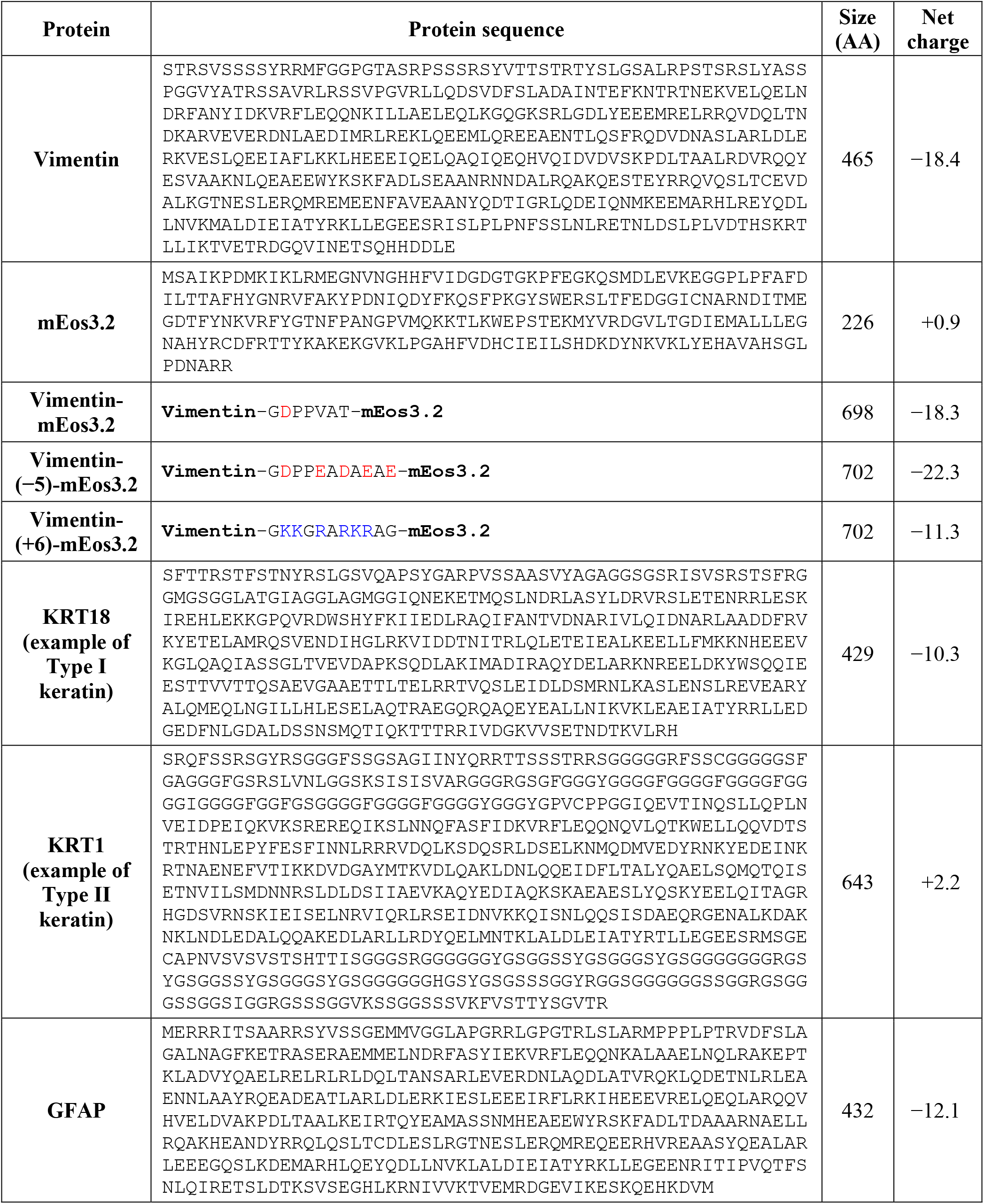
List of protein amino acid (AA) sequences and estimated net charges at pH = 7.3 per Protein Calculator v3.4 (http://protcalc.sourceforge.net

**Figure S1.**
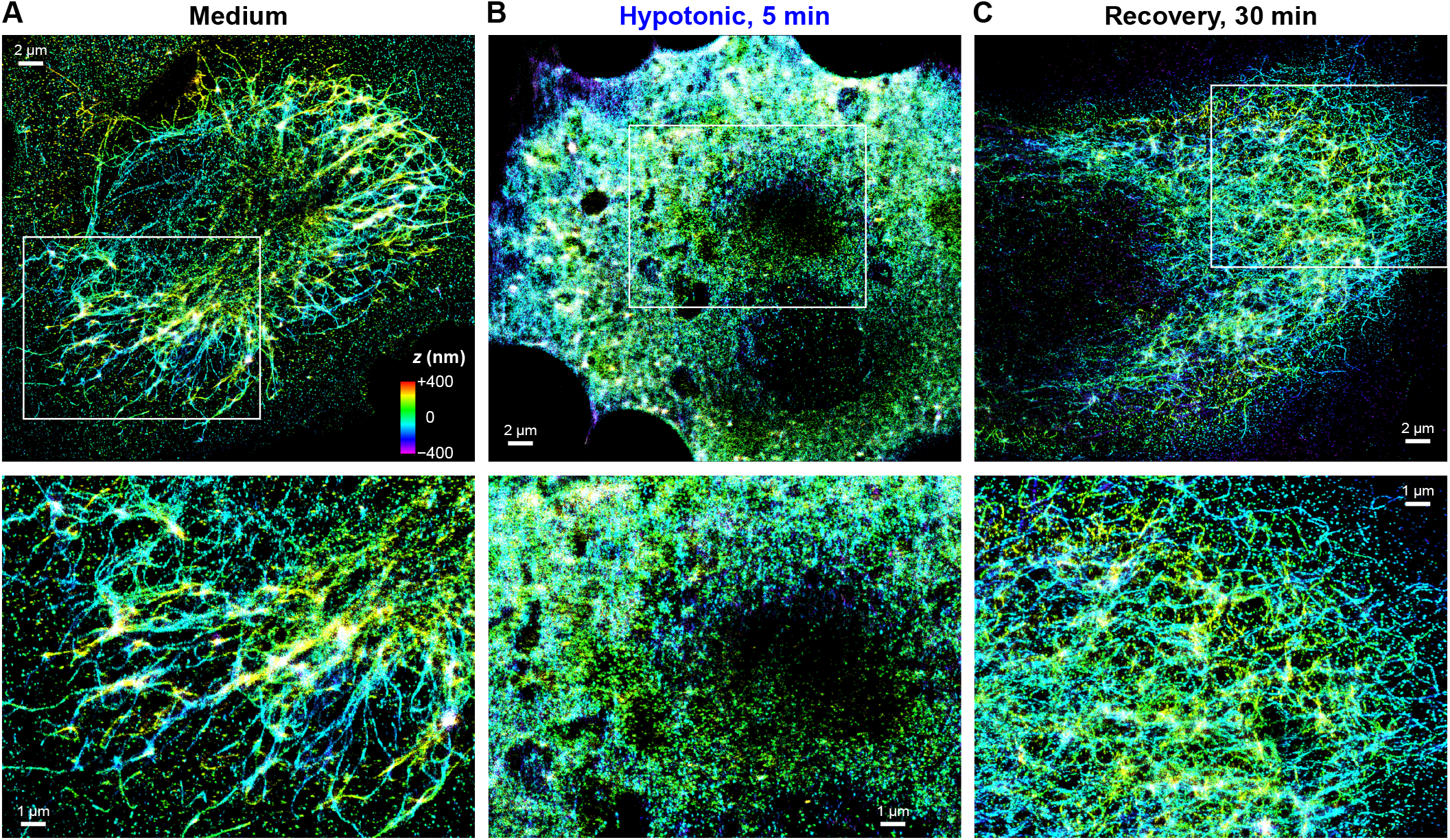
Representative 3D-STORM images of immunolabeled vimentin in COS-7 cells. (**A**) An untreated cell. The bottom figure is a zoom-in of the boxed region in the top figure. (**B**) Another cell, fixed after hypotonic treatment with water for 5 min. The zoom-in image shows the complete disassembly of filaments. (**C**) Another cell, after hypotonic treatment with water for 5 min, but then allowed to recover for 30 min in the regular cell culture medium in the incubator, before being fixed and labeled. Color encodes axial (depth) position, according to the color scale shown in (A).

**Figure S2.**
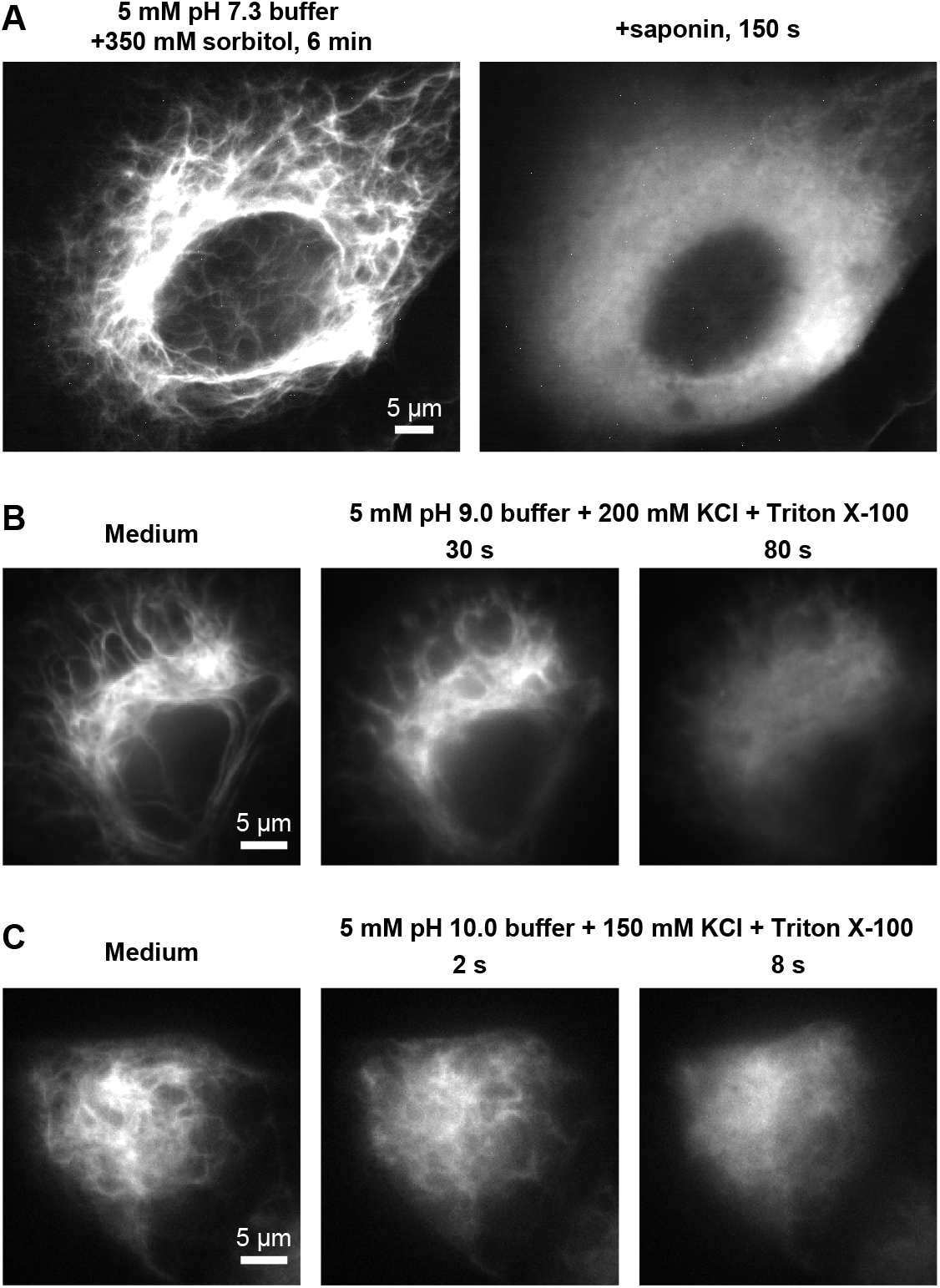
Additional fluorescence micrographs of vimentin-mEos3.2 in COS-7 cells after permeabilization under different ionic strengths and pHs. (**A**) Fluorescence micrographs of a cell (**i**) 6 min in a 5 mM phosphate buffer (pH = 7.3) with 350 mM sorbitol added and (**ii**) 150 s after next adding 50 µg/mL saponin into the medium. (**B**) Fluorescence micrographs of another cell before and after replacing the cell medium with a 5 mM CAPSO buffer (pH = 9.0) with the addition of 200 mM KCl and 0.2% Triton X-100, at 30 s and 80 s. (**C**) Fluorescence micrographs for another cell before and after replacing the cell medium with a 5 mM CAPSO buffer (pH = 10.0) with the addition of 150 mM KCl and 0.2% Triton X-100, at 2 s and 8 s.

**Movie S1**. Consecutive time series of the two-color live-cell data in **Fig. 4B**, shown as overlaid and separate color channels for vimentin-mCherry (magenta) and keratin-mEmerald (green). Time 0 corresponds to when the medium is changed to the 5 mM phosphate buffer. Scale bar: 5 μm.

## Notes

### Competing Interest Statement

The authors have declared no competing interest.

